# Modelling cancer immunomodulation using epithelial organoid cultures

**DOI:** 10.1101/377655

**Authors:** Yotam E. Bar-Ephraim, Kai Kretzschmar, Priyanca Asra, Evelien de Jongh, Kim E. Boonekamp, Jarno Drost, Joost van Gorp, Apollo Pronk, Niels Smakman, Inez J. Gan, Zsolt Sebestyén, Jürgen Kuball, Robert G.J. Vries, Hans Clevers

## Abstract

Here we utilize organoid technology to study immune-cancer interactions and assess immunomodulation by colorectal cancer (CRC). Transcriptional profiling and flow cytometry revealed that organoids maintain differential expression of immunomodulatory molecules present in primary tumours. Finally, we established a method to model antigen-specific epithelial cell killing and cancer immunomodulation in vitro using CRC organoids co-cultured with cytotoxic T cells (CTLs).

---

CRC is among the most common cancers worldwide^1^. While early CRC stages are highly treatable by surgical removal, later stages are usually incurable^2^. CRC arises through a multi-step process from small lesions of the epithelium of the large intestine. These lesions grow into adenomas with low grade dysplasia that progress into high grade dysplasia, eventually giving rise to infiltrating carcinomas^3^. Genetic mutations in signalling pathways such as the canonical Wnt signalling are the molecular basis of CRC^4^. However, the interaction of the tumour with its microenvironment is another critical hallmark^5^. Cancer cells remodel their microenvironment (e.g. fibroblasts, the vasculature and immune cells) to support tumour growth^6^. Infiltrating immune cells (ICs) such as CTLs or macrophages play a crucial role by generating different immune responses such as anti-tumour cytotoxicity (the former) or tumour-promoting chronic inflammation (the latter)^7^. As such, escape from the surveilling immune system has been recognised as one of the hallmarks of cancer^5^. Cancer cells undergo a process called immunoediting and silence anti-tumour responses, for example, by preventing T-cell activation through stimulation of inhibitory cell surface receptors such as CTL-associated antigen (CTLA)-4 or programmed death (PD)1^6,8^. Overcoming this active immunomodulation by tumour cells has become a major therapeutic target^9^. However, tumour heterogeneity, such as differential CTL infiltration or differential expression of immune inhibiting factors, could influence therapeutic efficiency of anti-tumour drugs by mediating drug resistance^6^. Developing *ex vivo* model systems to characterise the communication of the tumour with its environment is therefore of great importance. Organoid cultures grown from different epithelial tissues serve as an excellent tool to study tissue homeostasis and disease^10^. Furthermore, organoid biobanks of multiple epithelial organ systems have been established and tumour-derived organoids have successfully been used as platforms for screenings of different drugs to predict patient response^11^. Here we describe the establishment of a method to model antigen-specific epithelial-cell killing and cancer immunomodulation and *in vitro* using CRC organoids co-cultured with CTLs.

We first assessed whether CRC organoids expressed immunomodulatory molecules in established long-term expanded cultures. To this end, we compared gene expression of T-cell-specific immunomodulators in CRC organoids to the expression levels found in normal colon organoids using a transcriptome dataset generated using our ‘living organoid biobank’ of CRC patients^12^. On average, transcription of genes associated with T-cell stimulation such as *TNFSF4* or *TNFSF9* was not altered in CRC organoids compared to normal colon organoids (Fig. 1a). However, expression of human leukocyte antigen (HLA) genes *HLA-A* and *HLA-C*, encoding major histocompatibility complex class (MHC)-I molecules that present antigens to T cells, were significantly downregulated in CRC organoids (Fig. 1a), a well-described phenomenon found in cancers^13^. Expression of genes associated with inhibition of T-cell function was either significantly upregulated such as *BTLA*, significantly downregulated such as *CD80*, *CD86* or LGALS9 or not altered at all such as *CD274* (encoding PD-L1), *PDCD1LG2* (encoding PD-L2) (Fig. 1a). When assessing expression levels of immunomodulatory molecules on individual organoids, CRC organoids largely clustered together showing heterogeneous down regulation of *HLA-A*, *HLA-C* and *LGALS9* compared to healthy colon organoids (Fig. 1b). However, expression of immunoinhibitory genes *CD274* and *PDCD1LG2*, for instance, was highly upregulated in some CRC organoids in comparison to the matched normal colon organoid cultures, reflecting previously reported preservation of tumour heterogeneity in organoids^12^ (Fig. 1b). These molecular signatures provide a basis for further investigation of tumour immunogenicity and its association with other characteristics of the tumour.

**Fig 1.**
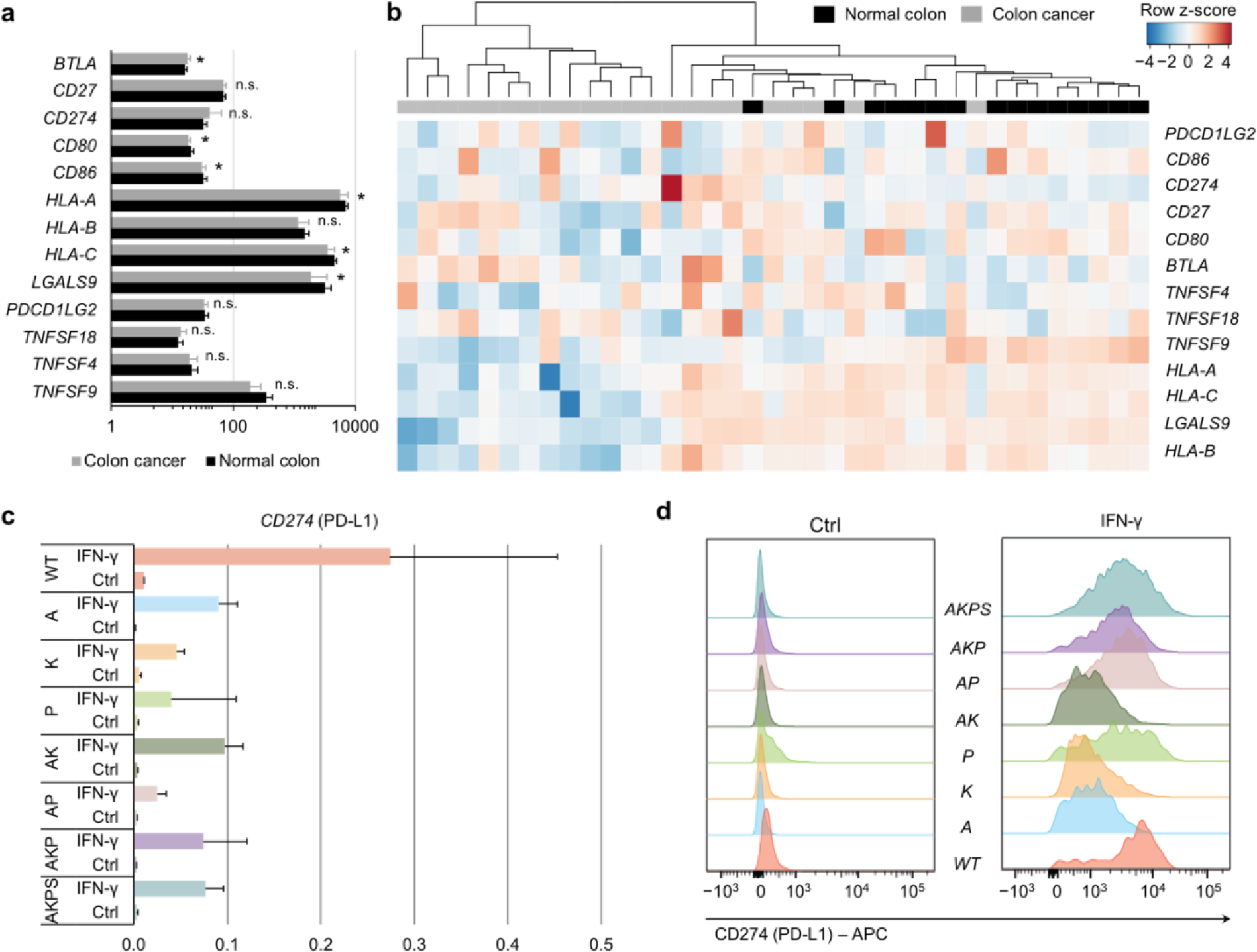
CRC organoids express immunomodulatory molecules. **a**,**b**, Normal colon and CRC organoid lines were generated in a patient-specific manner and RNA was extracted and analysed using Affymetrix single transcript microarrays. Average gene expression of different immunomodulators in normal colon and CRC organoid lines; n.s., non-significant; *, *p* < 0.05 (**a**). Hierarchical clustering of the individual normal colon and CRC organoid lines in the ‘living biobank’ displaying gene expression of selected immunomodulators. Color gradients represent z valued of each row (gene transcripts). **c-d**, Human colon organoid lines genetically engineered to carry one or more mutations found in CRCs. Expression levels of CD274 (PD-L1) in organoid lines (n = 2) at steady state (Ctrl) and upon stimulation with 20 ng/mL recombinant human IFN-γ assessed by quantitative PCR (**c**) and flow cytometry (**d**). A, *APC*^KO/KO^; N.D., not detected; K, *KRAS*^G12D/+^: P, *P53*^KO/KO^; S, *SMAD4*^KO/KO^, WT, wild-type.

Four of the most commonly mutated genes in CRC are *APC*, *P53*, *KRAS* and *SMAD4*, reflecting the stepwise progression of the normal intestinal epithelium into a metastatic carcinoma^14^. Introduction of these cancer mutations into human intestinal organoid cultures using clustered regularly interspaced short palindromic repeats (CRISPR)/Cas9 demonstrated that this process can be mimicked *in vitro* and upon xenotransplantation into mice^15,16^. Using colon organoids carrying one or more of these cancer mutations, we investigated whether up-regulation of PD-L1 was associated with a certain mutational status. Additionally, we exposed mutant organoids and their wild-type control organoid line to interferon (IFN)-γ, which is secreted by T cells and can trigger increased expression of immunomodulatory molecules such as PD-L1^17^. Subsequently, we assessed PD-L1 expression by quantitative polymerase chain reaction (qPCR) and flow cytometry (Fig. 1 c,d). In the absence of IFN-γ, organoids carrying triple (*APC*^KO/KO^, *P53*^KO/KO^, *KRAS*^G12D/+^) and quadruple mutations (*APC*^KO/KO^, *P53*^KO/KO^, *KRAS*^G12D/+^ and *SMAD4*^KO/KO^) showed lower *CD274* gene expression in comparison to control wild-type organoids (Fig. 1 c). Overall, PD-L1 expression was low in untreated organoid lines (Fig. 1 c,d). However, PD-L1 expression was dramatically upregulated in IFN-γ-treated organoids both on transcript and protein level (Fig. 1 c,d). These data demonstrate that CRC organoids express immunomodulators and that this expression is regulated in a similar way as previously shown for tissue *in vivo*.

We next aimed at establishing a co-culture system for CRC organoids and CTLs to model antigen-specific killing of tumour cells *in vitro*. For this, we used αβ T cells carrying a transgenic T-cell receptor (TCR) recognizing an HLA-A2-restricted Wilms tumour (WT)1-derived peptide^18,19^. We first screened CRC organoids from the ‘living biobank’^12^ as well as newly generated CRC organoids for HLA-A2 expression using flow cytometry. We found three CRC organoid lines that showed partial downregulation of HLA-A2 (Supplementary Fig. 1a). We were able to purify HLA-A2^+^ and HLA-A2^-^ CRC organoids and successfully established cultures from both populations (Fig. 2b). We confirmed stable MHC-I downregulation in HLA-A2^−^ CRC organoids, as IFN-γ stimulation did not trigger HLA-A2 re-expression (Supplementary Fig. 1b). Next, we pulsed these CRC organoid lines with WT1 peptide and, subsequently, co-cultured them for 48 hours with peptide-specific T cells. Following co-culture, we found that HLA-A2^−^ CRC organoids did survive irrespective of whether pulsed with the peptide or not (Fig. 2c). However, only the HLA-A2^+^ CRC organoids without prior peptide incubation survived co-culture (Fig. 2c). Peptide-pulsed HLA-A2^+^ CRC organoids were effectively killed by the peptide-specific T cells providing a proof-of-principle that organoids can be utilised to study anti-tumour response by cytotoxic T cells *in vitro*.

**Fig 2.**
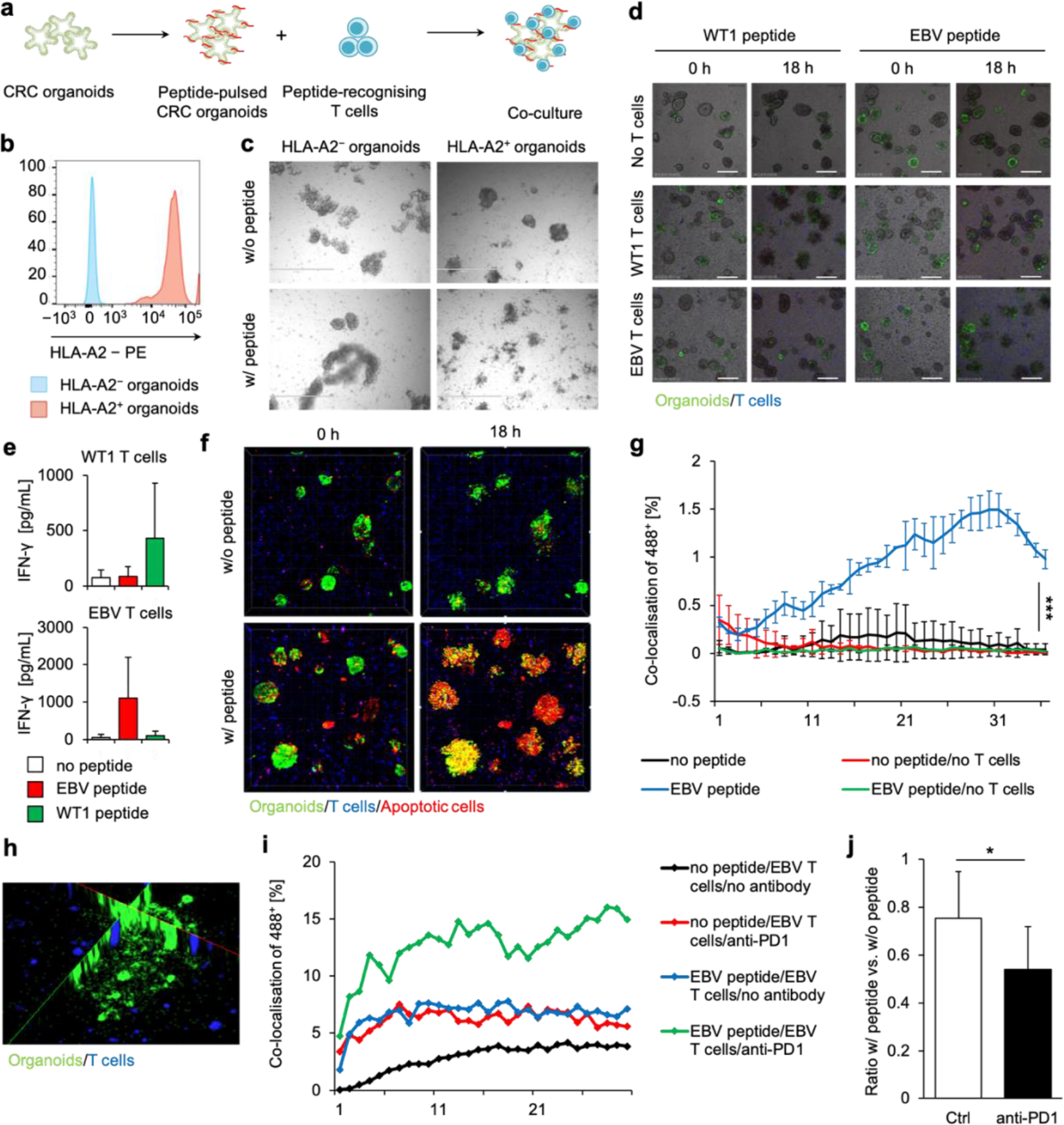
CRC organoids as tools for assessment of antigen specific killing by CD8^+^ T cells. **a**, Experimental scheme. **b**, Flow cytometry analysis of HLA-A2 expression in cloned HLA-A2^+^ and HLA-A2^-^ lines. **c**, Brightfield images of CRC organoids co-cultured with WT1 peptide-specific T-cell receptor-specific transgenic T cells for 48 hours; scale bars: 1 mm. **d**, Images showing peptide-pulsed HLA-A2^+^ CRC organoids at the beginning and end of co-culture with indicated peptide-specific T cells; scale bars: 70 μm. **e**, IFN-γ production by WT1 (top) and EBV (bottom) peptide-specific T cells as measured by ELISA of supernatants collected after 18-hour co-culture with HLA-A2^+^ CRC organoids pulsed with indicated peptides. **f**, Live-cell imaging stills of an 18-hour co-culture experiment with EBV peptide-pulsed HLA-A2^+^ CRC organoids co-cultured with an EBV-specific T-cell clone. **g**, Quantification of CRC organoid killing by specific T cells. Graphs are representative of multiple repeated experiments with either EBV peptide and EBV T-cell-or WT1 peptide and WT1 T-cell co-cultures. **h**, Representative projection image of T cells (blue) infiltrating a peptide-pulsed CRC organoid as recorded during the live-cell imaging experiments. **i**, Quantification of killing of IFN-γ treated CRC organoids by specific T cells in either presence or absence of a blocking antibody against PD-1. Graphs are representative of multiple repeated experiments with either EBV peptide and EBV T-cell- or WT1 peptide and WT1 T-cell co-cultures. **j**, Quantification cell viability after 18 hours co-cultures of either peptide pulsed or non-pulsed HLA-A2^+^ organoids with antigen specific T cells. Graphs represent ratio between peptide-pulsed and non-peptide pulsed conditions.

To further confirm antigen-specificity in our ‘killing’ assay system, we improved our co-culture method by transfecting HLA-A2^+^ CRC organoids with a construct expressing mNeonGreen-tagged histone H2B and staining T cells with CellTracker violet to allow for long-term tracking of both cell types (Methods). We then pulsed HLA-A2^+^ CRC organoids with either the WT1 peptide or with an EBV-derived peptide (Methods) and co-cultured the organoids with T cells carrying either a WT1- or an EBV-specific TCR. Here, only organoids pulsed with the cognate peptide were efficiently killed by the T cells (Fig. 2d, Supplementary Movies 1 and 2). Testing for IFN-γ production by the T cells in the co-culture using enzyme-linked immunosorbent assay (ELISA) confirmed antigen-specific organoid killing by the T cells (Fig. 2e). In order to better follow the kinetics of the organoid killing, we applied a fluorescent dye (NucRed Dead 647; Methods), which specifically stains apoptotic cells, and performed live confocal imaging on the co-culture (Fig. 2f, Supplementary Movies 1 and 2). We then quantified organoid killing by assessing co-localisation of NucRed Dead dye with H2B-mNeonGreen (Methods). Significant co-localisation of both labels and, hence, organoid killing, was only observed when peptide-pulsed HLA-A2^+^ CRC organoids were co-cultured with the respective peptide-specific T cells (Fig. 2g). Furthermore, T cells infiltrating into the epithelium of the organoids could be readily detected in this co-culture condition (Fig. 2h). Finally, we investigated whether using this co-culture system modulation of the immune response to immunosuppressive tumours can be modelled. Indeed, addition of a blocking antibody against PD-1 (αPD-1) enhanced tumour killing and IFN-γ production in PD-L1 expressing IFN-γ stimulated organoids (Fig. 2i,j). This was not observed when organoids were not IFN-γ stimulated and, hence, did not express PD-1. In conclusion, T cells efficiently killed co-cultured CRC organoids in an antigen-specific manner. In addition, T-cell inhibition and subsequent relief of this inhibition using αPD-1 could be modelled.

Here we demonstrate that epithelial organoids can be used to faithfully recapitulate the interaction between tumour tissue and the immune system. Also, using our co-culture assay, we set a first step in rebuilding the tumour microenvironment *in vitro*. Further addition of other components of this microenvironment (such as fibroblasts, natural killer cells, myeloid-derived suppressor cells, B cells) may shed light on the complex interactions between the different cell types leading to immune evasion of the tumour. Lastly, in line with a recent publication utilising cancer organoids as a scaffold for T-cell expansion^20^, this co-culture system can be used as a tool for drug-screens testing applicability of certain immunotherapies, for instance, chimeric antigen receptor (CAR)- or TCR transgenic T cells, antibody-dependent cell-mediated cytotoxicity (ADCC) or antibody-dependent cellular phagocytosis (ADCP) inducing antibodies directed at the tumour, to different tumours and different patients.

## Methods

### Human material and informed consent

Colonic tissues (both normal colon and tumour tissue) were obtained from the Departments of Surgery and Pathology of the Diakonessenhuis hospital, Utrecht, the Netherlands. All patients included in this study were diagnosed with CRC. Informed consent was signed by all included patients. Collection of tissue was approved by the medical ethical committee (METC) of the Diakonessenhuis hospital, in agreement with the declaration of Helsinki and according to Dutch and European Union legislation.

### Organoid generation and cultures

Epithelial organoid lines were derived from healthy colon or tumor tissue as previously described^12,21^. In brief, healthy colonic crypts were isolated by digestion of the colonic mucosa in chelation solution (5.6 mM Na_2_HPO_4_, 8.0 mM KH_2_PO_4_, 96.2 mM NaCl, 1.6 mM KCl, 43.4 mM Sucrose, and 54.9 mM *D*-Sorbitol, Sigma) supplemented with dithiotreitol (0.5 mM, Sigma) and EDTA (2 mM, in-house), for 30 minutes at 4°C. Colon crypts were subsequently plated in basement membrane extract (BME; Cultrex PC BME RGF type 2, Amsbio) and organoids were grown in human intestinal stem cell medium (HISC), which is composed of Advanced Dulbecco’s modified Eagle medium/F12 supplemented penicillin/streptomycin, 10 mM HEPES and Glutamax (all Gibco, Thermo Fisher Scientific) with 50% Wnt3a conditioned medium (in-house), 20% R-Spondin1 conditioned medium (in-house), 10% Noggin conditioned medium (in-house), 1 x B27, 1,25 mM n-acetyl cysteine, 10 mM nicotinamide, 50 ng/mL human EGF, 10 nM Gastrin, 500 nM A83-01, 3 μM SB202190, 10 nM prostaglandine E2 and 100 μg/mL Primocin (Invivogen). Tumor specimens were digested to single cells in collagenase II (1 mg/mL, Gibco, Thermo Scientific), supplemented with hyaluronidase (10 μg/mL) and LY27632 (10 μM) for 30 minutes at 37°C while shaking. Single tumor cells were plated in BME and organoids were cultured in HICS minus Wnt conditioned medium and supplemented with 10 μM LY27632 at 37°C.

### Organoid transfection

CRC organoids were dissociated into small clumps using TrypLE and then transduced with H2B-mNeonGreen (pLV-H2B-mNeonGreen-ires-Puro), as previously described^22^.

### T cells

Generation of αβ T cells carrying a transgenic TCR recognizing an HLA-A2-restricted WT1-derived peptide were described elsewhere^18^. Briefly, TCRα and β chains were cloned from raised tetramer positive T cell clones. Subsequently, CD8^+^ αβ TCR T cells were transduced using retroviral supernatant from Phoenix-Ampho packaging cells that were transfected with gag-pol, env, and pBullet retroviral constructs containing the cloned TCR genes.

### Organoid-T cell co-culture and live-cell imaging

Organoids stably transfected with H2B-mNeonGreen were split and digested a 5 to 7 days prior to co-culture and seeded at a density of 5000 cells per 10 μL of BME (25,000 cells per well in a 12-well cell culture plate). Two days prior to co-culture, T cells were starved from IL-2. One day prior to co-culture, organoids were stimulated with IFN-γ at indicated concentrations.

Prior to co-culturing, T cells were stained with Cell Proliferation Dye eFluor 450 (eBioscience) according to the manufacturer’s instructions. Organoids were pulsed with TCR-specific peptide (ProImmune) for 2 hours at 37°C prior to co-culture. Organoids and T cells were harvested and taken up in T cell medium, supplemented with 10% BME, 100 IU/ mL IL-2 and NucRed Dead 647 (Thermo Fischer). Where indicated, anti-PD1 blocking antibodies (2 μg/mL) were added to the co-culture. Cells were plated in glass-bottom 96-well plates and co-cultures were imaged using an SP8X confocal microscope (Leica).

### Flow cytometry

APC-labelled pentamers to the EBV-derived, HLA-2:02 restricted peptide FLYALALLL (ProImmune) where used to sort pentamer^+^ CD8^+^ CD3^+^ T cells from PBMCs isolated from buffycoats from healthy individuals. Cells were sorted as single cells into 96-well plates using a BD FACS Aria (BD Biosciences) cytometer. For flow cytometry, the following antibodies were used (all anti-human): CD8^−^ PE (clone RPA-T8), CD45^−^ PerCP-Cy5.5 (2D1), CD274 (PD-L1)^−^ APC (MIH1) (all BD Biosciences), CD279 (PD-1)^−^ PE (EH12.2H7, Biolegend), HLA-A2^−^ PE (BB7.2, Santa Cruz).

### Quantitative polymerase chain reaction (qPCR)

For qPCR analysis, RNA was isolated from organoids using the RNAeasy kit (QIAGEN) according to the manufacturer’s protocol. PCR analysis was performed using the SYBR Green Reagent (Biorad). PCR reactions were performed in duplicate with a standard curve for every primer. Primers were designed using the NCBI primer design tool. Primers used in this study: *GAPDH* forward (GTC GGA GTC AAC GGA TT), *GAPDH* reverse (AAG CTT CCC GTT CTC AG), *HPRT* forward (GGC GTC GTG ATT AGT GAT), *HPRT* reverse (AGG GCT ACA ATG TGA TGG), *CD274* forward (TGC AGG GCA TTC CAG AAA GAT), *CD274* reverse (CCG TGA CAG TAA ATG CGT TCAG).

### Transcriptional profiling

Microarray analysis of biobank organoids was performed as described elsewhere^12^.

### Enzyme linked immunosorbent assays (ELISA)

Culture supernatants were kept at −20°C and ELISA was performed for indicated cytokines using ELISA MA Standard (Biolegend) according to manufacturer’s protocol.

### Cell viability assay

Cell viability after co-cultures was assessed using CellTiter-Glo Luminescent cell viability assay (Promega), according to manufacturer’s protocol.

### Image analysis

Image analysis was done using Imaris software package (Bitplane). In brief, threshold for positive staining was set on negative controls. A co-localization channel was made for H2B-neon and NucRed Dead 647 signals. Cell death was quantified as percentage of H2B-mNeonGreen^+^ voxels co-localising with NucRed Dead signal.

### Bioinformatics analysis

Bioinformatics analysis of normalised gene-expression data from microarray experiments12 was performed using standard packages (*i.e.* gplots) in R version 3.4.0 (R Foundation, https://www.r-project.org) and RStudio version 1.0.143 (https://www.rstudio.com).

### Statistical analysis

All experiments were repeated at least three times unless otherwise indicated. All data were shown as mean ± SEM. Statistical significance was analysed by either ANOVA or two-tailed Student’s *t*-test using either Graphpad Prism 6 or Microsoft Excel 2010.

## Acknowledgements

We thank T. Aarts, S. van den Brink, A. Cleven, T. Mizutani, M. Schiffler, M. van de Wetering for technical assistance and experimental advice, A. de Graaff and the Hubrecht Imaging Facility for help with microscopy, S. van der Elst, R. van den Linden and T. Poplonski for help with flow cytometry. This work was supported by the European Research Council (Advanced Grant ERC-AdG 67013-Organoid to H.C.) and a VENI grant from the Netherlands Organisation for Scientific Research (NWO-ZonMW, 016.166.140 to K.K.). This work is part of the Oncode Institute, which is partly funded by the Dutch Cancer Society. K.K. is a long-term fellow of the Human Frontier Science Program Organization (HFSPO, LT771/2015) and was a long-term fellow of the European Molecular Biology Organisation (EMBO, ALTF 839-2014).

## Author contributions

Y.B.E. and K.K. designed, performed and analysed the experiments and wrote the manuscript. Y.B.E. performed image analysis. K.K. performed bioinformatics analysis. P.A., E.d.J. and K.E.B. assisted with experiments. J.D. generated cancer gene-mutant organoid lines. J.v.G. isolated tumour and normal tissue from resected material. A.P. and N.S. performed surgery. I.J.G., Z.S. and J.K. provided WT1 peptide and WT1 peptide-specific transgenic TCR αβ T cells. R.G.J.V. organised tissue collection. K.K. and H.C. acquired funding. H.C. supervised the project and wrote the manuscript. All of the authors commented on the manuscript.

